# Use of a plasmid containing a dual gene reporter system to assess the cell hydrophobicity of *Listeria monocytogenes*

**DOI:** 10.64898/2026.06.06.730570

**Authors:** Ogueri Nwaiwu, Catherine Rees

## Abstract

*Listeria monocytogenes* causes listeriosis in humans and animals and contaminates prepared food by attaching to food processing environments. Therefore, closer monitoring of how the organism adheres to surfaces will help identify ways to prevent it from colonising food-processing environments. To develop new attachment assays, clinical and environmental strains of *L. monocytogenes* were transformed by inserting a plasmid containing *lux, gfp* reporter genes and an erythromycin-resistant gene into the parent cells. Transformed cells were grown for 48 hours on brain heart infusion agar plates containing 1-5µg/ml of erythromycin, after which the cells were viewed under a molecular light imager and luminometer. Fluorescent cells containing the *gfp, lux*, and erythromycin-resistant genes were visible, whereas control cells without the plasmid were not. Transformation efficiency was highest with the environmental strains, and subsequent growth and hydrophobicity tests carried out with the transformed cells in different growth conditions showed that they were able to attach well to solvents when compared to the parent cells. However, the growth rate of the transformed cells was poor, indicating a disruption of cell metabolism. Results show the possibility of real-time monitoring of how cells attach to different surfaces and could lead to a better understanding of the initial colonisation of a surface by the organism.

## 1.0 INTRODUCTION

*Listeria monocytogenes* is a Gram-positive non-spore-forming bacterium and a facultative intracellular pathogen but exists as a gut commensal and is also found commonly in soil, surface water, sewage, vegetation, and food processing plants. Since 1926, when Listeria monocytogenes was first described, it has continued to show worldwide prevalence associated with serious diseases in humans and a wide variety of animals. On brain heart infusion agar, the colonies are small, round, and milky white on reflected light. The ability of the microbe to grow in a wide variety of temperatures makes it an effective food pathogen that can be found in unprocessed food of animal origin, such as fish, milk, meat, and poultry. and, in contaminated vegetables (Nwaiwu, 2016) or salad. The organism can multiply rapidly in aerobic or microaerophilic environments under conditions of pH between 6 and 9.

*L. monocytogenes* can cause Listeriosis, a serious food-borne bacterial infection that can occur due to the organism’s ability to resist gastric and bile acids, colonise the intestinal lumen, cross the intestinal barrier, survive intracellularly in the bloodstream, evade immune responses, and cross the placental and blood-brain barriers (Disson et al., 2025).

The organism is a constant threat to the food industry and public health because we are exposed by our nutritional habits (Mantovam et al, 2025). An analysis of literature data by Díaz-Martínez et al. (2025) on potential associations between specific antibiotic resistance patterns in the pathogen and food categories found that food type had an influence on the antibiotic resistance profiles of *L. monocytogenes*, with meat and vegetables being the food categories exhibiting the most prevalent profiles. This resistance may be aided by the acquisition of plasmids from the environment (Nwaiwu, 2022).

There is a need to develop new methods of monitoring the organism to have a better understanding of its behaviour in different environments. To this end, the use of bioprotective (Fliss et al, 2025) and endolysin-based (Kin et al., 2025) strategies has been proposed. Another way to monitor and quantify the growth of the organisms in real time may be by using bioluminescence, and Hou et al. (2025) have suggested the use of luminescent nanomaterials. Some bacteria are known to convert energy into visible light, which allows them to produce light and glow in the dark. Daunert et al (2000) explained that the bacteria luciferase catalyses the luminescence reaction in luminous bacteria to produce visible light. Light production catalysed by the luciferase in bacteria plays a role in the protection of the cells against oxidative stress (Szpilewska *et al*, 2003). The genes and the gene products necessary for bacterial bioluminescence have been identified (Engerbrecht and Silverman, 1984).

Bacterial cells can also be monitored *In situ* using green fluorescent protein (GFP). GFP is known as a photoprotein found in the jellyfish Aequorea victoria. It has been pointed out by Chudakov *et al* (2005) that fluorescent labelling is of paramount importance to biological studies, and researchers have extensively utilised GFPs for designing fluorescence biosensors due to their intrinsic fluorescence, high stability (Tian et al., 2023), An advantage of GFP as a reporter protein is its auto-fluorescence, and therefore its use does not require the addition of any cofactors or exogenous substances to produce light. The development of various fluorescent proteins has opened novel applications for *In vivo* fluorescent labelling in protein expression studies (Griesbeck, 2004). The ability of *L. monocytogenes* to express GFP has been demonstrated (Jian et al, 2005), and it has been used (Freitag and Jacobs, 1999) to examine the organism’s intracellular gene expression. Furthermore, it can be used to develop new methods of analysis for ecological and industrial applications (Ramesh and Prasastha, 2025).

There is a need for more knowledge and understanding of the mechanisms of *L. monocytogenes* adaptation to environmental stress factors, which will lead to the development of new methods of pathogen control in the food industry (Osek et al., 2022). Hence, this study was carried out to explore the improvement of real-time monitoring of *L. monocytogenes* during attachment to surfaces.

## 2.0 Materials and methods

### 2.1 Plasmid, bacteria strains, media, and cell hydrophobicity

The plasmid and *L*.*monocytogenes* strains used were obtained from the University of Nottingham, United Kingdom culture collection. The plasmid (Punk3008) used for electroporation harboured *lux, gfp* and erythromycin-resistant genes. Cell hydrophobicity of *Listeria* cells grown in three different growth media, namely rich brain heart infusion (BHI), minimal D10, was prepared as shown in a previous study (Nwaiwu et al, 2021). The D10 media supplemented with 10% duck meat extract (DJ+D10) was also prepared. The factory isolates used are described in that study, and the reference cells used for comparison were from well-known characterised strains ATCC 23074 (4b) and Lm 10403S (1/2a).

### 2.2. Preparation of competent L. monocytogenes cells and electroporation with a plasmid

Overnight cultures (10ml) were grown in BHI/0.5M sucrose solution, after which 100ml BHI/ sucrose was inoculated with the overnight culture to 0.5 OD. This was grown with shaking (Gallenkamp) at 37 °C to A600=0.2. Penicillin (Sigma) was added to a final concentration of 10/ml, following which the cells were grown for another 2 hours to OD of 0.35 – 0.4.

Cells were pelleted at 5000g under a temperature of 4 deg for 10 minutes and re-suspended in an equal volume (100ml) of HEPES(1mM, pH 7.0) - Sucrose (0.5M) solution. Cells were pelleted again under the same conditions and re-suspended in half volume (50ml) HEPES -Sucrose solution. This was repeated after which the cells were drained carefully and re-suspended in 1/400 HEPES - Sucrose solution (.25ml). Plasmid DNA(Punk3008) was dialysed simultaneously with a 0.025 μm filter membrane (Millipore) for 30 minutes and added to 100 μL of competent cells, following which the mixture was introduced into an electroporation cuvette (Geneflow).

Electroporation was carried out with Gene Pulser (Bio-Rad) at a pulse of 2.0, resistance 200 Ohms and a capacitance of 250 µF, after which the cells were resuspended in 900µl of BHI – Sucrose solution before expressing for 2 hours. Transformed *Listeria* colonies were obtained by plating out the expressed cells on BHI supplemented with 1µg/ml of erythromycin. Incubation was carried out for 48 hours after which colonies that emerged were counted and sub-cultured for single colonies into higher concentration (5µg/ml) of BHI plates containing erythromycin. The number of transformants and the transformation efficiency were calculated.

### 2.3 Measurement of fluorescence and luminescence expression of parent and transformed cells of L. monocytogenes

This was carried out to quantify fluorescence and luminescence of the transformed cells. Transformed and parent cells were grown in 5 mL of BHI overnight, after which the cells were pelleted at 5000g. Before pelleting the cells, the OD of the cells was recorded. The cells were washed twice in 1ml of PBS and then pelleted at 13000 g, after which re-suspension was carried out with 200µl of PBS. The cells were then loaded in a microtiter plate before measuring luminescence with a light meter (Victor^™^). The transformed cells were checked for evidence of fluorescence by visualising under the NightOWL imager (Berthold Technologies). Three independent tests were performed.

### 2.4 Growth of Listeria monocytogenes in broth of D10, BHI and D10 supplemented with duck meat extract (1:10)

For each broth, overnight cells (5 mL) were pelleted at 5000g. The pelleted cells were introduced into 50ml broth in 250ml conical flasks and grown with shaking at 30°C and 150 rpm. Growth was measured at 620nM by taking OD readings at intervals of 30 mins or 1 hour with a Cecil 20 spectrophotometer. The same procedure was repeated for transformed cells of *Listeria monocytogenes* in addition to measurement of their luminescence as growth progressed.

Luminescence during growth was carried out by placing 1.5ml of broth culture in a scintillating tube (Meridian), after which luminescence was measured (Turner 20 Luminometer). The generation time and growth rate of the parent and transformed cells were calculated using the standard equations

A)The Specific growth rate (µ)

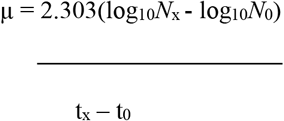

B)The generation (doubling) time (g)

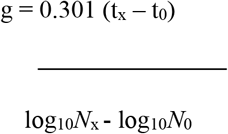

## 3.0 Results and Discussion

### 3.1 Electroporation of competent Listeria monocytogenes cells with Plasmid

Two clinical reference strains, Lm10403S (1/2a) and Lm 23074 (4b) and three environmental strains were used. The environmental strain with the same serotypes as the reference strains were chosen to determine how the environmental and clinical isolates attach to surfaces and determine if serotype would have any effect on attachment. Cells of *L. monocytogenes* were transformed with a plasmid containing the *lux, gfp* reporter genes and the selective erythromycin resistance gene. Transformed cells of *L. monocytogenes* grown on BHI plates containing 1µg of erythromycin were able to grow after 48 hours.

When the cells were viewed under the molecular light imager, fluorescent cells containing the *gfp, lux* and erythromycin resistant gene were seen (Fig 1). The cells were able to grow when further sub-culturing was performed on higher concentration (5µg/ml) of erythromycin plates (Fig 1b). Transformation efficiency was highest with Lm101 (Table 1).

**Table 1.**
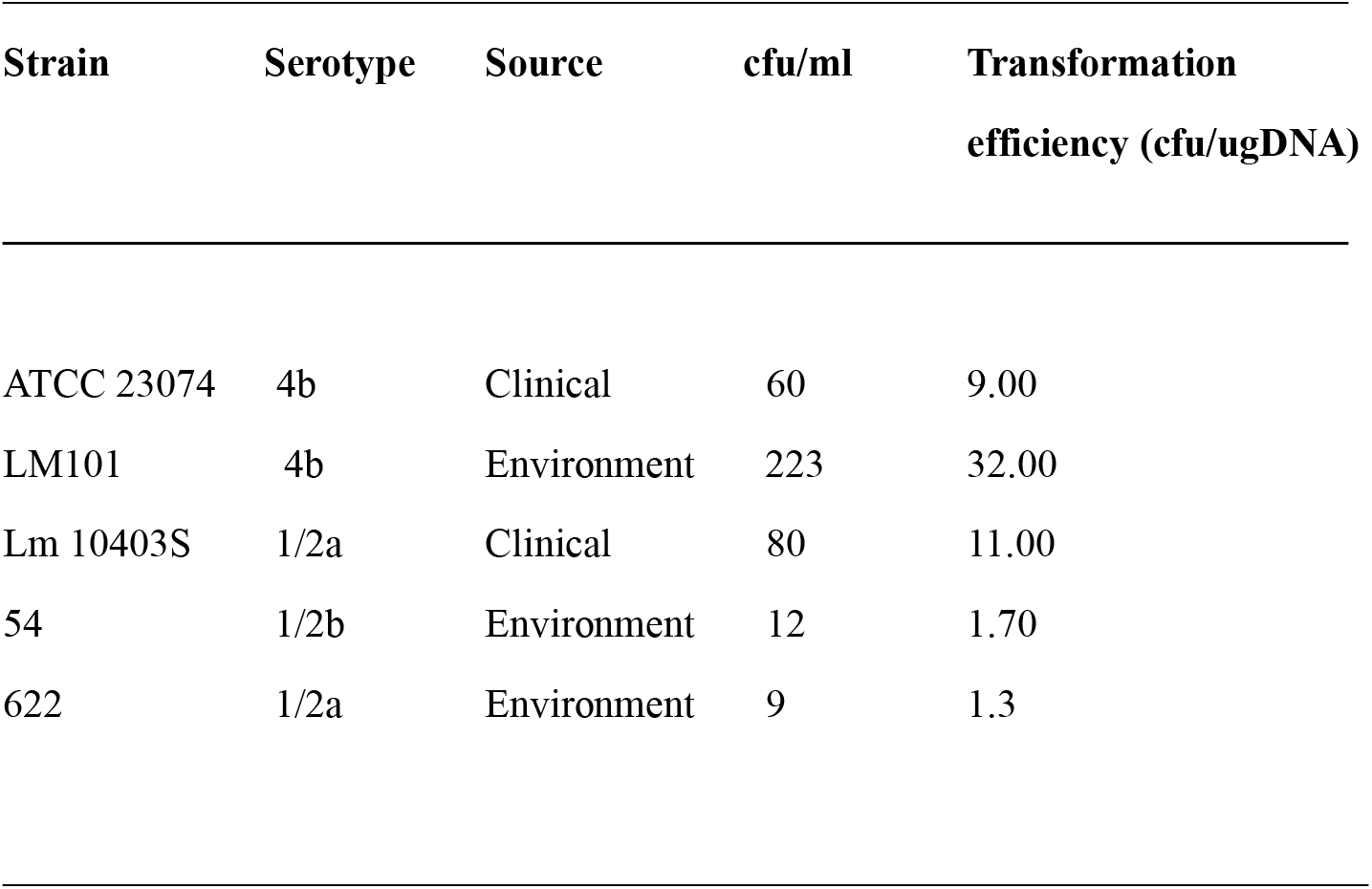
Transformation efficiency for electroporation performed with approximately 0.7 μg of plasmid. The environment strain Lm101 showed the highest transformation efficiency.

| Strain | Serotype | Source | cfu/ml | Transformation<br>efficiency (cfu/ugDNA) |
| --- | --- | --- | --- | --- |
| ATCC 23074 | 4b | Clinical | 60 | 9.00 |
| LM101 | 4b | Environment | 223 | 32.00 |
| Lm 10403S | 1/2a | Clinical | 80 | 11.00 |
| 54 | 1/2b | Environment | 12 | 1.70 |
| 622 | 1/2a | Environment | 9 | 1.3 |

**Fig 1.**
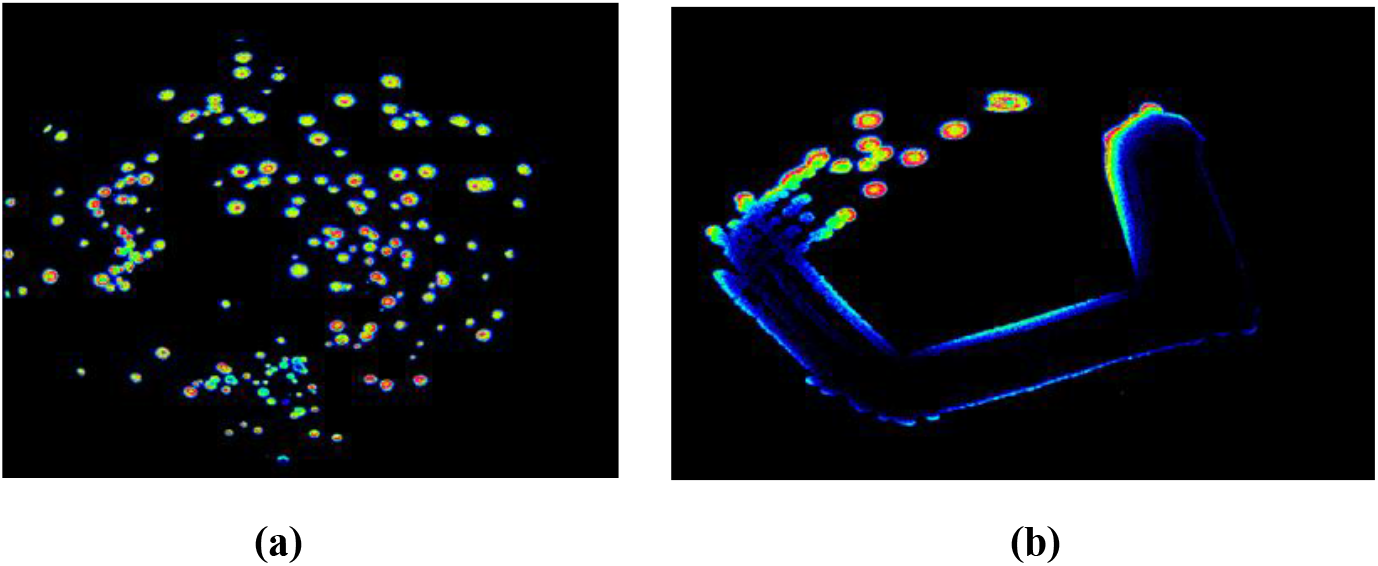
Transformed cells of *L. monocytogenes* grown on BHI media. Cells were visible with media supplemented with 1 µg (Fig 1a) and 5 µg (Fig 1b) of erythromycin.

### 3.2 Measurement of fluorescence and luminescence expression of parent and transformed cells of L. monocytogenes

After electroporation, the quantity of fluorescence and luminescence was measured to determine the suitability of the transformed cells for use as biosensors. Even though all the cells had an optical density above 1.5 (Fig 2a) before washing with PBS, the environmental strains Lm101 and Lm54 had more fluorescence (Fig 2b) and luminescence (Fig 2c) than the clinical strains.

**Fig 2a.**
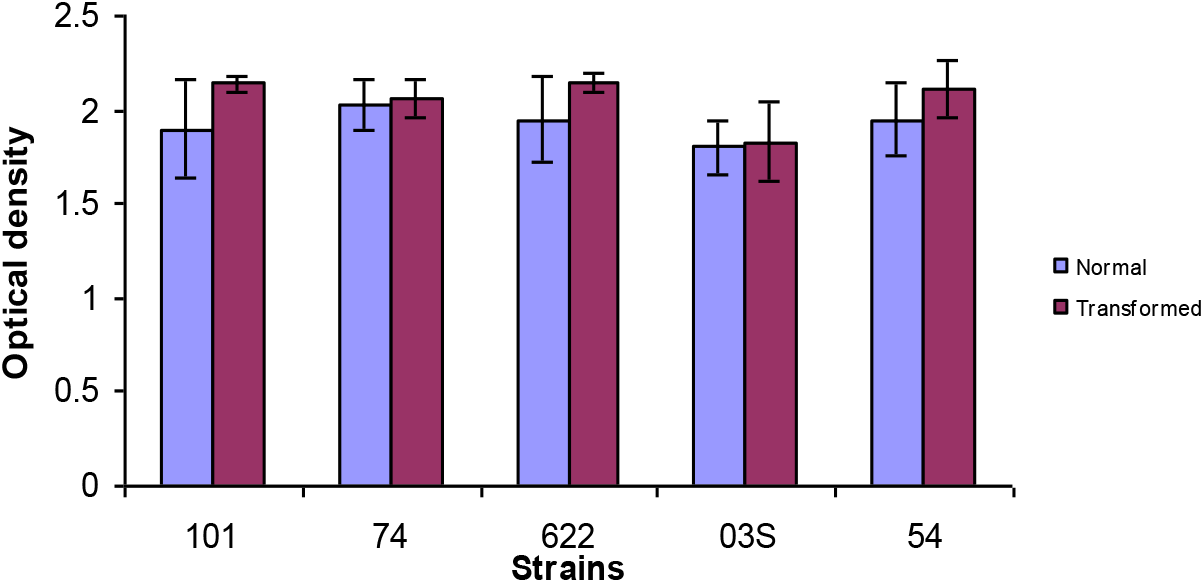
Optical density of transformed and parent cells before bioluminescence and fluorescence measurements.

**Fig 2b.**
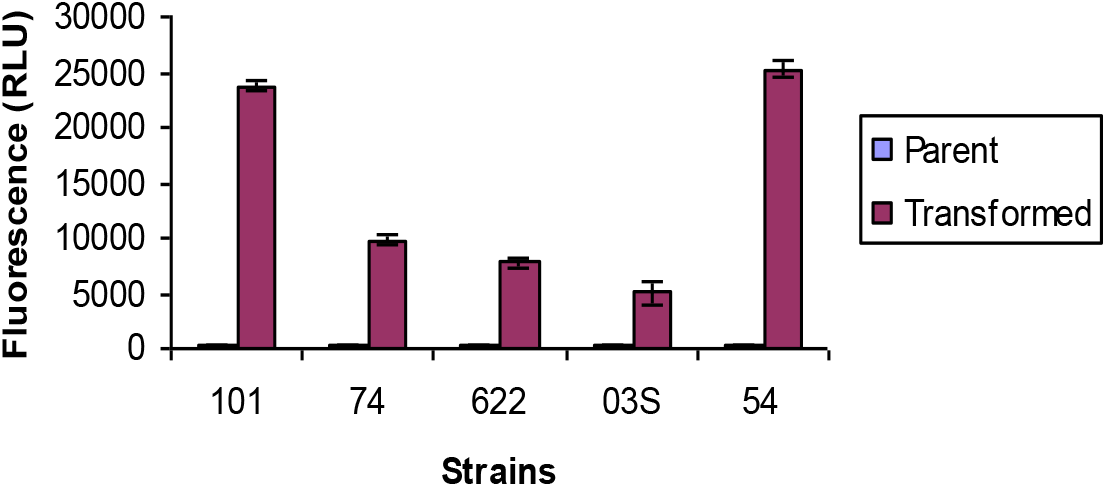
Fluorescence measurement of transformed cells.

**Fig 2c.**
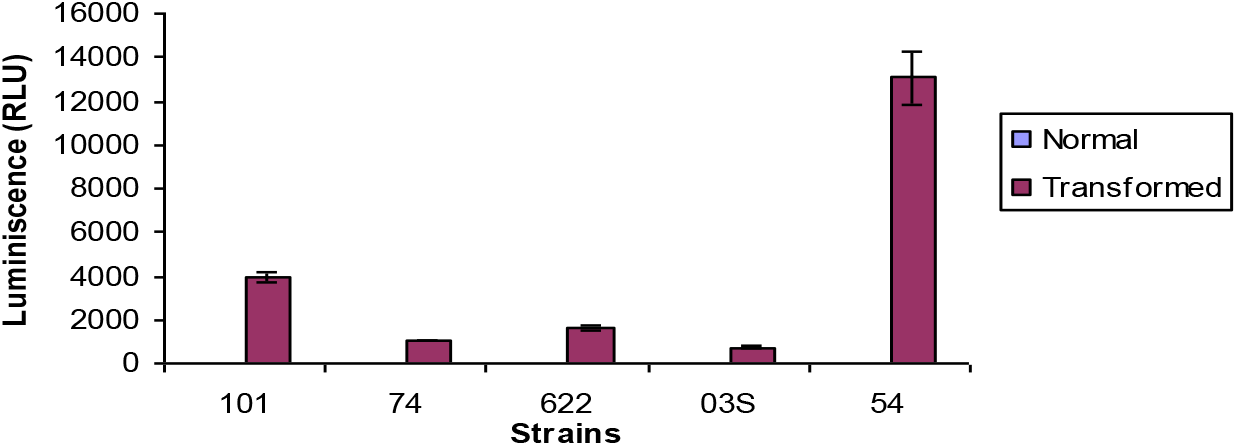
Luminescence measurement of transformed cells.

### 3.3 Growth and luminescence of Listeria monocytogenes in broth of D10, BHI and D10 supplemented with duck meat (1:10)

Duck juice was extracted from duck meat fillets obtained from a supermarket. Physiological analysis of the growth of the parent and transformed cells in a minimal nutrient environment was performed by growing the cells in a rich BHI medium, a minimal D10 medium and D10 medium supplemented with duck juice. For the transformed cells, luminescence during growth was also measured.

The Growth and luminescence of *Listeria monocytogenes* in broth of D10, BHI and D10 supplemented with duck meat (1:10) is shown in figures 3-7. In all the strains tested, growth was highest in BHI, followed by duck meat / D10 mix and lastly D10. Luminescence increased with growth and decreased after 4-5 hours. In BHI, it increased for the first 100 minutes and then levelled off. This suggests that the cell metabolism of the transformed cells is highest during the first 4 hours of growth, after which it decreases.

**Fig 3.**
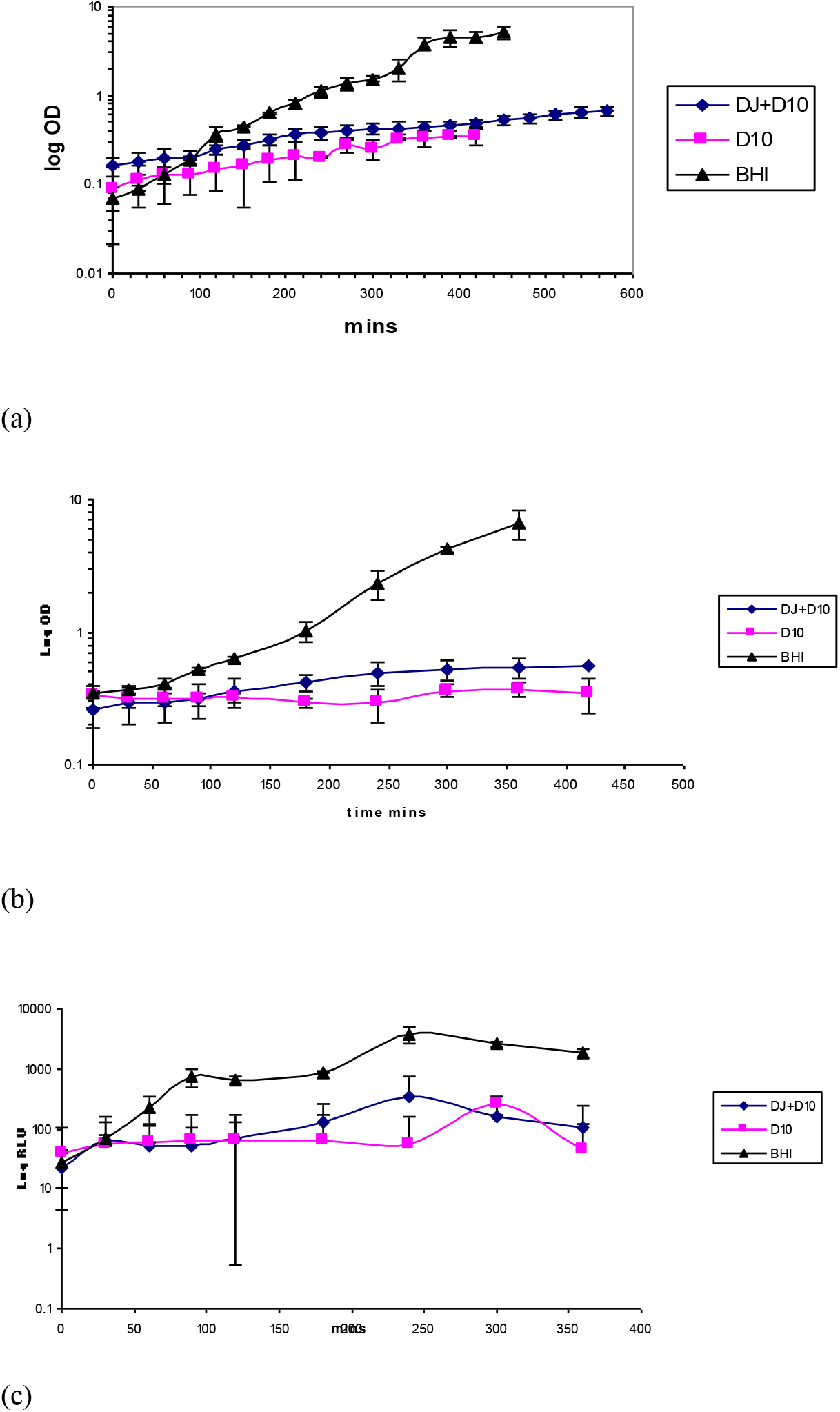
Growth of Lm23074 in broth of D10, BHI and D10 supplemented with duck meat (1:10). Growth for the parent cells (a) was significantly higher in BHI followed by D10 supplemented with duck meat and D10 media having the least growth. The growth was significantly higher in the transformed cells (b) Luminescence increased with growth and then started to diminish after 4-5 hours (c)

**Fig 4.**
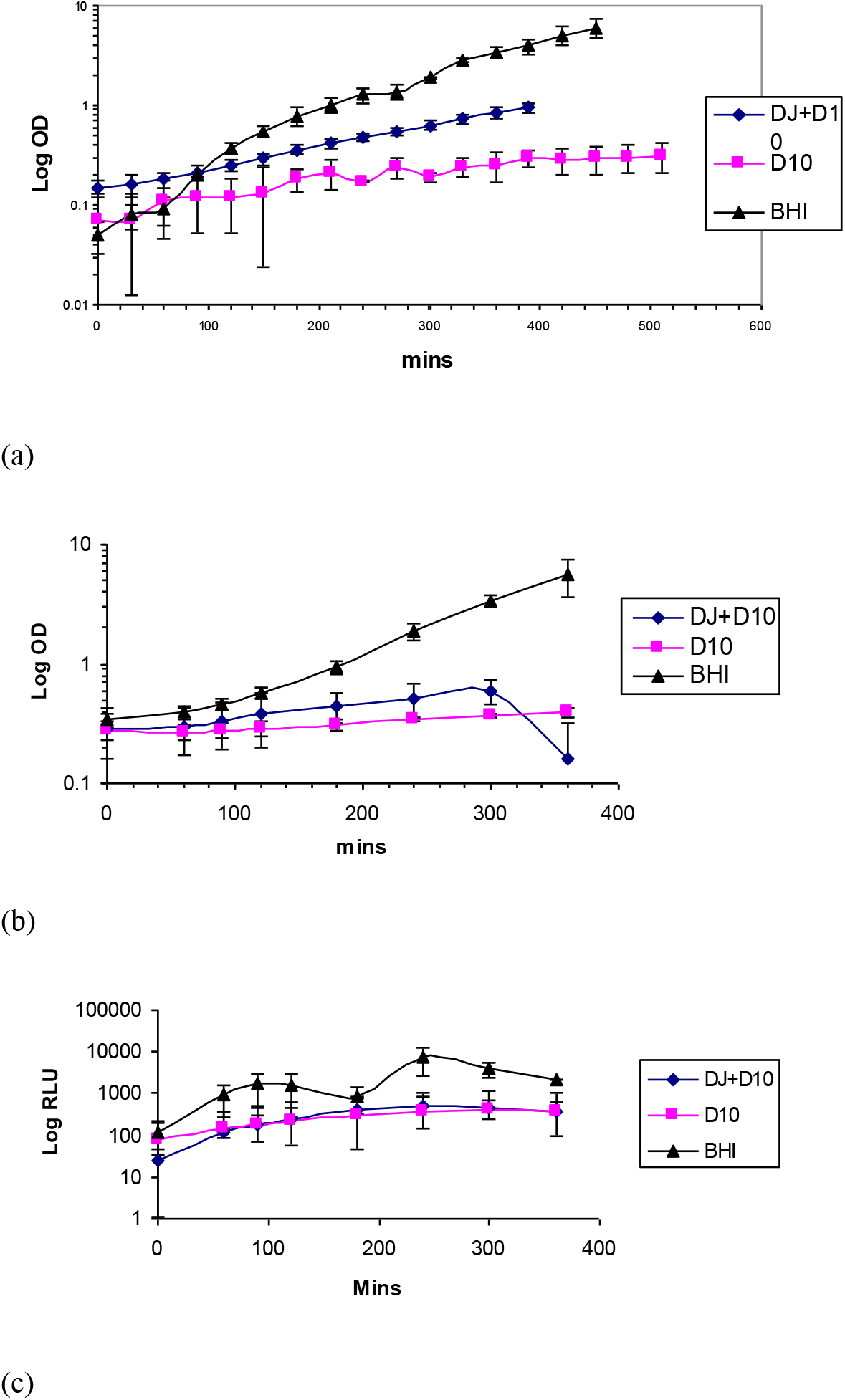
Growth of Lm101 in broth of D10, BHI and D10 supplemented with duck meat (1:10). Also, growth for the parent cells (a) was significantly higher in BHI The growth was significantly higher in the transformed cells (b). Luminescence increased with growth and then started to diminish after 4-5 hours (c).

**Fig 5.**
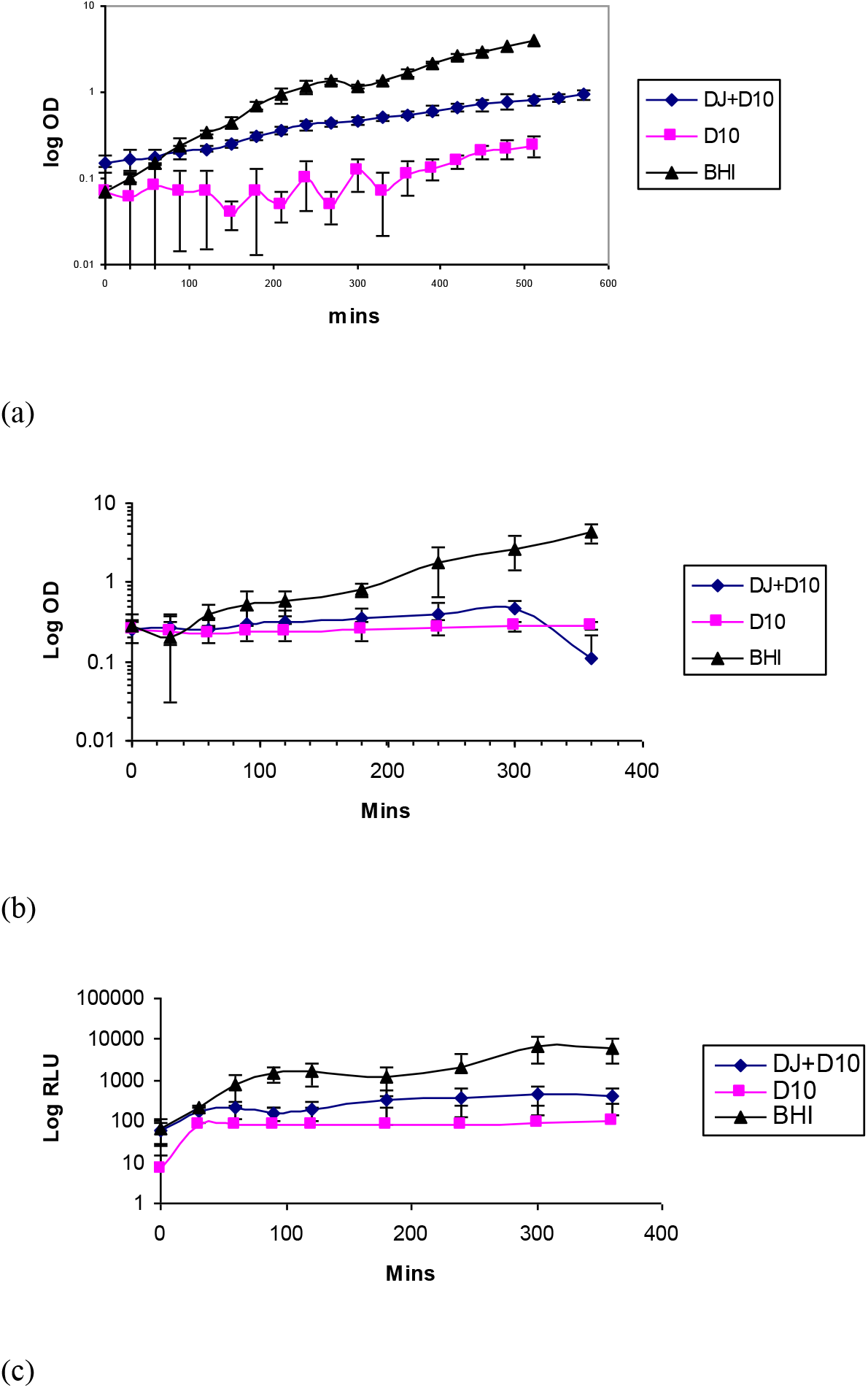
Growth of Lm54 in broth of D10, BHI and D10 supplemented with duck meat (1:10).

**Fig 6.**
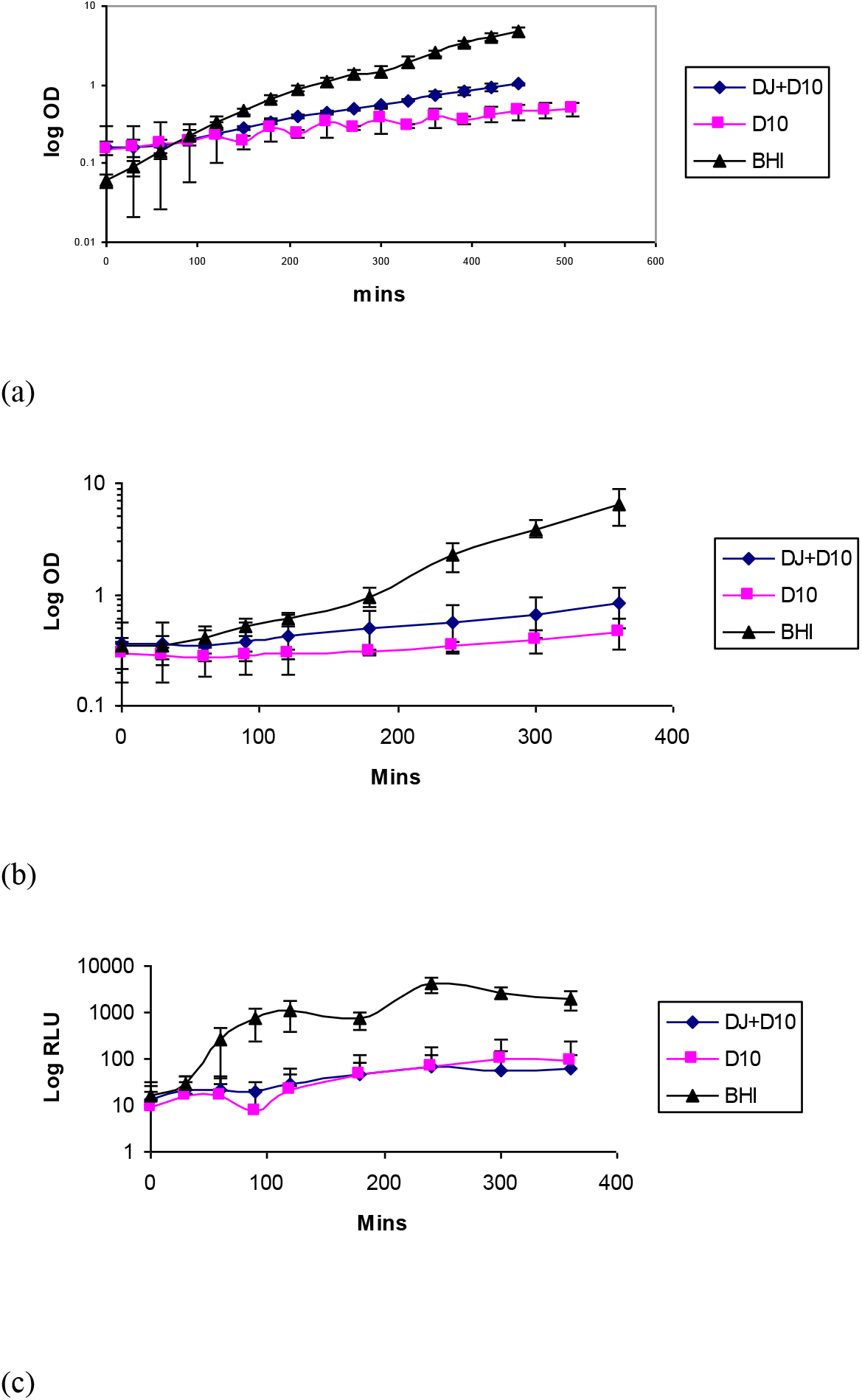
Growth of Lm622 in broth of D10, BHI and D10 supplemented with duck meat (1:10).

**Fig 7.**
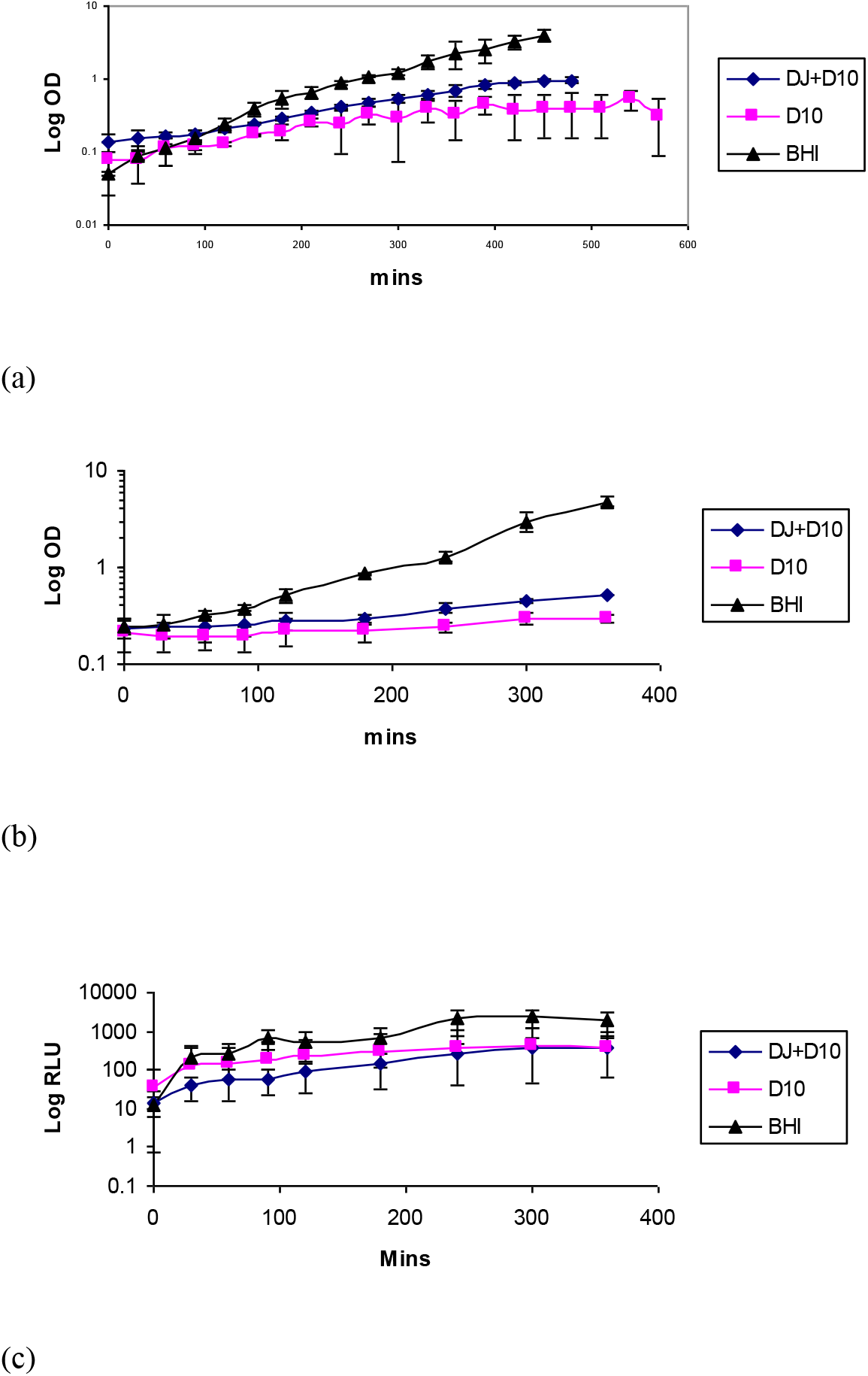
Growth of Lm 10403S in broth of D10, BHI and D10 supplemented with duck meat (1:10).

The growth rate and generation time analysis (Table 2) showed that the parent cells had a lower generation time and higher growth rate than the transformed cells. In the parent cells, it was observed that the environmental serotype 4b (Lm 101) had a lower generation time and higher growth rate in BHI and D10 media with duck meat supplement than the clinical serotype 4b (Lm23074).

**Table 2.** Generation time and growth rate of parent and transformed cells of *Listeria monocytogenes*.

| **Parent cells** |  |  |  |  |  |  |
| --- | --- | --- | --- | --- | --- | --- |
| **Strain** | **Generation time (g)** |  |  | **Growth rate ($\mu$)** |  |  |
|  | **BHI** | **DJ+D10** | **D10** | **BHI** | **DJ+D10** | **D10** |
| Lm 10403S | 16.44 | 133.78 | 143.96 | 2.53 | 0.31 | .29 |
| 622 | 12.88 | 128.43 | 265.59 | 3.23 | 0.32 | 0.16 |
| Lm23074 | 30.43 | 361.20 | 265.59 | 1.37 | 0.12 | 0.16 |
| 101 | 15.33 | 143.33 | 301.00 | 2.71 | 0.29 | .014 |
| 54 | 17.67 | 252.00 | 209.91 | 2.35 | 0.17 | 0.20 |
| **Transformed cells** |  |  |  |  |  |  |
| Lm 10403S | 60.84 | 2257.50 | 4267.00 | 0.68 | 0.02 | 0.01 |
| 622 | 58.01 | 2883.26 | 6086.89 | 0.72 | 0.01 | 0.01 |
| Lm23074 | 97.83 | 259.19 | 416.77 | 0.43 | 0.16 | 0.10 |
| 101 | 96.91 | 186.62 | 537.50 | 0.43 | 0.22 | 0.08 |
| 54 | 42.00 | 3763.00 | 8772.00 | 0.98 | 0.01 | 0.00 |

However, the clinical serotype had a lower generation time and higher growth rate in the D10 media than the environmental strain. This same pattern was observed when clinical serotype 1/2a (10403S) was compared with environmental serotype 1/2a (Lm622). It is possible that the clinical strains used can withstand nutritional stress.

## 4.0 Conclusions

The *L. monocytogenes* strain 4b had a higher transformation efficiency, suggesting it may have greater potential to acquire plasmids from the environment. The reporter genes used may be useful for visualising cells subjected to hydrophobicity in both rich and minimal media. The generation time and growth rate of transformed cells are lower than those of the parent cells, but growth improves with 10% duck juice supplementation.

## Notes

### Competing Interest Statement

The authors have declared no competing interest.

## REFERENCES

Chudakov, D.M., Lukyanov, S. and Lukyanov, K.A. (2005) Fluorescent proteins as a toolkit for in vivo imaging. Trends in Biotechnology 23, 605–613.

Díaz-Martínez, C., Bolívar, A., Mercanoglu Taban, B., Kanca, N., & Pérez-Rodríguez, F. (2025). Exploring the antibiotic resistance of Listeria monocytogenes in food environments -a review. Critical reviews in microbiology, 51(5), 731–754. 10.1080/1040841X.2024.2412007

Disson, O., Charlier, C., Pérot, P., Leclercq, A., Paz, R. N., Kathariou, S., Tsai, Y. H., & Lecuit, M. (2025). Listeriosis. Nature reviews. Disease primers, 11(1), 71. 10.1038/s41572-025-00654

Engebrecht, J. and Silverman, M. (2003) Identification of genes and gene products necessary for bacterial bioluminescence. Proceedings of the National Academy of Sciences 81, 4154–4158.

Fliss, O., Fliss, I., & Biron, E. (2025). Bioprotective Strategies to Control Listeria monocytogenes in Food Products and Processing Environments. International journal of molecular sciences, 26(21), 10481. 10.3390/ijms262110481

Freitag, N. E., & Jacobs, K. E. (1999). Examination of Listeria monocytogenes intracellular gene expression by using the green fluorescent protein of Aequorea victoria. Infection and immunity, 67(4), 1844–1852. 10.1128/IAI.67.4.1844-1852.1999

Griesbeck, O. (2004) Fluorescent proteins as sensors for cellular functions. Current Opinion in Neurobiology 14, 636–641.

Hou, S., Lin, T., Ding, Z., Cui, S., Cao, Y., Jiao, J., Kang, Q., & Du, X. (2025). Nanoparticles-based lateral flow immunochromatographic strip for detection of foodborne pathogen: A review. Food chemistry, 496(Pt 1), 146595. 10.1016/j.foodchem.2025.146595

Jiang, L. L., Song, H. H., Chen, X. Y., Ke, C. L., Xu, J. J., Chen, N., & Fang, W. H. (2005). Characterization of a mutant Listeria monocytogenes strain expressing green fluorescent protein. Acta biochimica et biophysica Sinica, 37(1), 19–24. 10.1093/abbs/37.1.19

Kim, D. W., Kong, M. S., & Koo, O. K. (2025). Endolysin-based biocontrol strategies against Listeria monocytogenes in food: A comprehensive review. Food science and biotechnology, 34(14), 3119–3125. 10.1007/s10068-025-01849-4

Liu, Dongyou (2006) Identification, subtyping, and virulence determination of Listeria monocytogenes as an important food pathogen. Journal of Medical Microbiology, 55, 645–659.

Mantovam, V. B., Dos Santos, D. F., Giola Junior, L. C., Landgraf, M., Pinto, U. M., & Todorov, S. D. (2025). Listeria monocytogenes, Salmonella spp., and Staphylococcus aureus: Threats to the Food Industry and Public Health. Foodborne pathogens and disease, 22(12), 809–824. 10.1089/fpd.2024.0124

Nwaiwu O. (2022). Comparative genome analysis of the first Listeria monocytogenes core genome multi-locus sequence types CT2050 AND CT2051 strains with their close relatives. AIMS microbiology, 8(1), 61–72. 10.3934/microbiol.2022006

Nwaiwu, O., Wong, L., Lad, M., Foster, T., MacNaughtan, W., & Rees, C. (2021). Properties of the Extracellular Polymeric Substance Layer from Minimally Grown Planktonic Cells of Listeria monocytogenes. Biomolecules, 11(2), 331. 10.3390/biom1102033

Nwaiwu, O. (2016). Molecular serotype and evolutionary lineage of Listeria monocytogenes isolated from different Nigerian food items. African Journal of Biotechnology, 15(17), 696–705.

Osek, J., Lachtara, B., & Wieczorek, K. (2022). Listeria monocytogenes -How This Pathogen Survives in Food-Production Environments?. Frontiers in microbiology, 13, 866462. 10.3389/fmicb.2022.866462

Ramesh, C., & Prasastha, V. R. (2025). Unlocking the potential of fluorescent microbes: Exploring their ecological and industrial applications. Biotechnology advances, 83, 108628. 10.1016/j.biotechadv.2025.108628

Tian, F., Xu, G., Zhou, S., Chen, S., & He, D. (2023). Principles and applications of green fluorescent protein-based biosensors: a mini-review. The Analyst, 148(13), 2882–2891. 10.1039/d3an00320e

